# CXCR1 and CXCR2 display receptor bias for shared chemokine agonists

**DOI:** 10.1101/2025.09.30.679682

**Authors:** Chanpreet Jassal, Joseph Strawn, Krishna Rajarathnam, Sudarshan Rajagopal

## Abstract

G protein–coupled receptors (GPCRs) mediate diverse signaling outputs through their proximal transducers: G proteins, GRKs, and β-arrestins. Although ligand bias at chemokine receptors (CKRs), where ligands for the same receptor display distinct signaling patterns, is well recognized, receptor bias, where the same agonist at different receptors yields distinct transducer engagement, remains poorly understood. We compared endogenous chemokine ligands (CXCL1, CXCL5, CXCL7, CXCL8) at the highly homologous CXCR1 and CXCR2 receptors using biosensor assays to measure Gαi activation, β-arrestin1/2 recruitment, GRK2/3/5/6 translocation, and receptor internalization. Our data reveal qualitatively different signaling patterns, most notably where CXCL1 acts as a G protein–biased partial agonist at CXCR1 but as a balanced full agonist at CXCR2. These signaling differences correlate with receptor internalization but not subcellular ERK activation patterns measured using compartment-specific biosensors. Collectively, our findings demonstrate receptor bias in CKR signaling, transducer activation, and compartmentalized kinase activation in translating chemokine identity into discrete functional outcomes.

**SIGNIFICANCE STATEMENT:** Chemokine ligand bias, where different ligands for the same receptor display different signaling patterns, is now well appreciated. However, there are only few examples of receptor bias, where the same agonist generates distinct signaling profiles at different receptors. Here we used biosensors and compartmental ERK biosensors, to show that CXCL1, CXCL5, CXCL7, and CXCL8 differentially engage G proteins, β-arrestins, and GRKs, at CXCR1 and CXCR2. This work provides mechanistic insight into CXCR1/CXCR2 signaling diversity.

## INTRODUCTION

G Protein-Coupled Receptors (GPCRs) are the most common class of receptors found throughout the human body, and are the target of approximately 35% of all U. S. Food and Drug Administration–approved drugs (Sriram and Insel, 2018). GPCRs mediate signaling by engaging many transducers, such as heterotrimeric G proteins, GPCR Kinases (GRKs), and β-arrestins (Eiger et al., 2022). Most ligands that bind GPCRs are thought to display “balanced” agonist activity, signaling through pathways mediated by β-arrestins and G proteins (Rajagopal et al., 2010b). However, it has been appreciated that some receptor-ligand systems selectively signal through some pathways over others in a ligand-, receptor-, or systems-dependent manner − a phenomenon referred to as “biased agonism” (Smith et al., 2018). Given the broad range of physiological processes regulated by GPCRs and the association of their dysfunction with numerous pathological conditions (Schoneberg et al., 2004), investigating biased signaling to enhance efficacy and reduce side-effect profiles of existing GPCR-based therapies offers significant promise.

Moreover, GPCRs with multiple endogenous agonists can display bias between those ligands (Smith et al., 2018). The human chemokine system represents a subfamily of GPCRs where endogenous bias is critical in modulating diverse cellular functions. It is composed of approximately 50 known endogenous chemokine ligands and 20 known chemokine receptors (CKRs), displaying a spectrum of selective and promiscuous interactions at both the ligand and receptor level resulting in biased responses (Eiger et al., 2021). Biased agonism, specifically ligand bias, in the chemokine system was first described at CCR7, where studies demonstrated selective signaling through either β-arrestins or G proteins (Byers et al., 2008; Kohout et al., 2004; Zidar et al., 2009). Later, it was appreciated that some CKRs such as ACKR3, also known as CXCR7, are β-arrestin-biased and signal exclusively through β-arrestin in the absence of G protein activation (Rajagopal et al., 2010a). However, the complexity of biased agonism at CKRs remains to be fully characterized. This is especially true of receptor bias, the ability of the same agonist to result in distinctly different signaling patterns depending on the receptor to which it binds (Smith et al., 2018).

C-X-C motif chemokine receptor 1 (CXCR1) and C-X-C motif chemokine receptor 2 (CXCR2) are the major CKRs expressed on neutrophils and mediate their recruitment to infection sites to trigger cytotoxic effects (Nasser et al., 2009) (Rajarathnam et al., 2019). CXCR1 and CXCR2 are highly homologous and share 77% amino acid identity with a majority of the divergent residues clustered in three regions: (i) the N-terminal segment before transmembrane domain (TMD) 1, (ii) the region from TMD4 to the end of the second extracellular loop (ECL2), and (iii) the C-terminal cytoplasmic tail (Ahuja et al., 1996; Murphy and Tiffany, 1991). While it is well established that CXCR1 and CXCR2 are high-affinity receptors for CXCL8, they also bind six additional chemokines (CXCL1–3, 5–8) with varying affinities (Boon et al., 2024; Boon et al., 2025). These chemokines share the characteristic Glu-Leu-Arg (ELR) motif at the N-terminus that is critical for activating both receptors. While studies have focused on ligand bias between these chemokines at either CXCR1 and CXCR2 (Boon et al., 2024; Boon et al., 2025; Rajagopal et al., 2013), little is known regarding the presence of receptor bias in the response of the same chemokine agonist between these two closely related receptors. Towards this goal, we characterized how biased signaling at CXCR1 and CXCR2 receptors and their endogenous chemokines, CXCL1, CXCL5, CXCL7, and CXCL8 influences unique intracellular signaling events through β-arrestins, G proteins, and GRK transducers.

## MATERIALS AND METHODS

### Generation of constructs

Construct cloning was performed using conventional techniques such as overlap cloning as previously described unless otherwise indicated (Bryksin and Matsumura, 2010).

### Bacterial strains

XL-10 Gold ultracompetent E. coli (Agilent) was used to express all constructs used in this manuscript (Table S1).

### Cell Culture and Transfection

Human Embryonic Kidney (WT HEK293T, β-arrestin 1/2 knockout) cells were grown in minimum essential media (MEM) supplemented with 10% fetal bovine serum (FBS) and 1% penicillin/streptomycin (P/S) at 37 °C and 5% CO2. β-arrestin 1/2 CRISPR knockout cells were provided by Stephane Laporte (McGill University, Montreal, Quebec, Canada) (Namkung et al., 2016). For BRET and luminescence-based assays transient transfections were performed using the polyethylenimine (PEI) transfection method. Cell culture media was replaced 30 min prior to transfection. Plasmid constructs were suspended in Opti-MEM (GIBCO) to a final volume of 100 μL. In a separate tube, 100 μL of PEI in Opti-MEM was prepared at a PEI:DNA ratio of 3:1 using a PEI stock concentration of 1mg/mL. After 5 minutes, PEI solution was added to the plasmid DNA, gently mixed, and allowed to incubate at room temperature (RT) for 30 minutes. The PEI:DNA mixture were then added to the cells.

### TRUPATH G-protein Activation Assays

HEK293T cells seeded in 6-well plates were transiently transfected using PEI with wildtype CXCR1 or CXCR2, Gαi1-RLuc8, Gγ9-GFP2, and Gβ3 at equal amounts (Olsen et al., 2020). 24 hours later, cells were plated onto clear bottom, white-walled, 96-well plates (Costar) at 100,000 cells/well in clear MEM supplemented with 1% FBS and 1% P/S. The following day, the media were aspirated, and cells were incubated at room temperature with 80 μL of coelenterazine 400a (5 μM final concentration) in Hanks’ balanced salt solution (HBSS) (Gibco) supplemented with 20 mM HEPES for 5 minutes. BRET signals were measured with a BioTek Synergy Neo2 plate reader at 37°C plates with a standard 400 nm emission filter and 510 nm long pass filter following addition with 20 μL of ligands(10 μM of chemokines). BRET ratios were calculated by dividing the 510 nm signal by the 400 nm signal. Net BRET values were calculated by subtracting the vehicle BRET ratio from the ligand-stimulated BRET ratio.

### EKAR BRET Assays

HEK293T cells seeded in 6-well plates were transiently transfected using PEI with wildtype CXCR1 or CXCR2, EKAR BRET biosensors tagged to different cellular locations, and dynamin K44A or pcDNA control. 24 hours later, cells were plated onto clear bottom, white-walled, 96-well plates (Costar) at 100,000 cells/well in clear MEM supplemented with 1% FBS and 1% P/S with or without 200 ng/mL pertussis toxin (PTX). The following day, the media were aspirated, and cells were incubated at room temperature with 80 μL of coelenterazine h (2.5 μM final concentration) in Hanks’ balanced salt solution (HBSS) supplemented with 20 mM HEPES for 5 minutes. BRET signals were measured with a BioTek Synergy Neo2 plate reader at 37°C using a 480nm wavelength filter (NLuc) and 530nm wavelength filter (mVenus). Three prereads were taken to quantify the baseline BRET signals before 20 μL of ligands were added (10 μM of chemokines). BRET ratios were calculated by dividing the 530 nm signal by the 480 nm signal. Net BRET values were calculated by subtracting the vehicle BRET ratio from the ligand-stimulated BRET ratio.

### β-arrestin Recruitment and Receptor Internalization BRET Assays

HEK293T and β-arrestin 1/2 knockout cells seeded in 6-well plates were transiently transfected using PEI with CXCR1/CXCR2 tagged with a C-terminal RLuc8, and β-arrestin1 or β-arrestin2 tagged with a C-terminal YFP for β-arrestin recruitment assays or Fyve-tagged mVenus for receptor internalization. 24 hours later, cells were plated onto clear bottom, white-walled, 96-well plates (Costar) at 100,000 cells/well in clear MEM supplemented with 1% FBS and 1% P/S with or without 200 ng/mL pertussis toxin (PTX). The following day, the media were aspirated, and cells were incubated at room temperature with 80 μL of coelenterazine h (2.5 μM final concentration) in Hanks’ balanced salt solution (HBSS) supplemented with 20 mM HEPES for 5 minutes. BRET signals were measured with a BioTek Synergy Neo2 plate reader at 37°C using a 480nm wavelength filter (NLuc) and 530nm wavelength filter (mVenus). Three prereads were taken to quantify the baseline BRET signals before 20 μL of ligands were added (10 μM of chemokines). BRET ratios were calculated by dividing the 530 nm signal by the 480 nm signal. Net BRET values were calculated by subtracting the vehicle BRET ratio from the ligand-stimulated BRET ratio.

### NanoBiT GRK Recruitment Assays

HEK293T cells seeded in 6-well plates were transiently transfected using PEI with wildtype CXCR1 or CXCR2, GRK2/GRK3/GRK5/GRK6 were tagged with a C-terminal LgBiT (Kawakami et al., 2022), and a CD8α-SmBiT (plasma membrane marker). 24 hours after transfection, cells were plated onto clear bottom, white-walled, 96-well plates (Costar) at 100,000 cells/well in clear MEM supplemented with 1% FBS and 1% penicillin/streptomycin with or without 200 ng/mL pertussis toxin (PTX). The following day, the media were aspirated, and cells were incubated at room temperature with 80 μL of coelenterazine h (2.5 μM final concentration) in HBSS Gibco supplemented with 20 mM HEPES for 5 minutes. Luminescence signals were read using a BioTek Synergy Neo2 plate reader at 37°C read without a wavelength-specific filter. Three prereads were taken to quantify the baseline luminescence before adding 20 μL of ligands at the appropriate concentrations. The average of the luminescence prereads was divided from each read following ligand addition to calculate a change in luminescence over baseline and then normalized to vehicle treatment.

### CXCR1 and CXCR2 Ligands

CXCL1, CXCL5, CXCL7, and CXCL8 were recombinantly expressed and purified as described previously (Ravindran et al., 2013). The chemokines were and diluted in Dulbecco’s phosphate-buffered solution (PBS) supplemented with 0.1% bovine serum albumin and aliquots were stored at −80°C until needed for use.

### Statistical Analyses

Statistical analyses were performed using GraphPad Prism 10 (GraphPad Software). Values are reported as mean ± SEM. Dose-response curves were fitted to a log agonist versus stimulus with three parameters (span, baseline, and *EC*_50_) with minimum baseline corrected to zero. When comparing ligands or treatment conditions in concentration-response assays, a two-way ANOVA of ligand and concentration was performed. For time course experiments, pairwise comparison using a one-way ANOVA followed by Tukey’s multiple comparison’s test was conducted on the AUC. An unpaired t-test was used for comparing cell lines effects on receptor internalization. P<0.05 was considered to be statistically significant. Further details of statistical analysis and replicates are reported in the figure legends. Bias plots were generated using normalized dose-response data for G protein activation, β-arrestin1 and β-arrestin2 recruitment, and receptor internalization and plotting responses at the same concentration of chemokine for each set of analyses.

## RESULTS

### Chemokines promote differential activation of G proteins and β-arrestins at CXCR1 and CXCR2

We first assessed G protein signaling activity at CXCR1 and CXCR2 with four endogenous chemokine ligands − CXCL1, CXCL5, CXCL7, and CXCL8. Using the TRUPATH bioluminescence resonance energy transfer (BRET) assay (Olsen et al., 2020) (**Fig. 1A**), we assessed CXCR1 and CXCR2 signaling by measuring heterotrimeric Gαi dissociation. Following chemokine stimulation of CXCR1, CXCL8 displayed the most Gαi activation, followed by CXCL7, CXCL1 and CXCL5 (**Fig. 1B**). In contrast, while every chemokine promoted Gαi activity at CXCR2, no differences in maximal response between ligands were observed (**Fig. 1C**). Given the role of β-arrestin2 in desensitizing G protein signaling, we next examined β-arrestin2 recruitment to CXCR1 and CXCR2 following chemokine stimulation by BRET (**Fig. 1D**). At CXCR1, all ligands recruited β-arrestin2 (**Fig. 1E**) with rank order of CXCL8 > CXCL7 > CXCL1 > CXCL5 (**Fig. 1F** and **G**). In contrast, at CXCR2, both CXCL8 and CXCL1 generated maximal responses, closely followed by CXCL7, and then CXCL5, which showed the lowest β-arrestin2 activity (Fig.1H-J). Similar trends were observed with β-arrestin1 recruitment for all chemokines at both CXCR1 and CXCR2 (**Supplemental Figure 1**).

**Figure 1.**
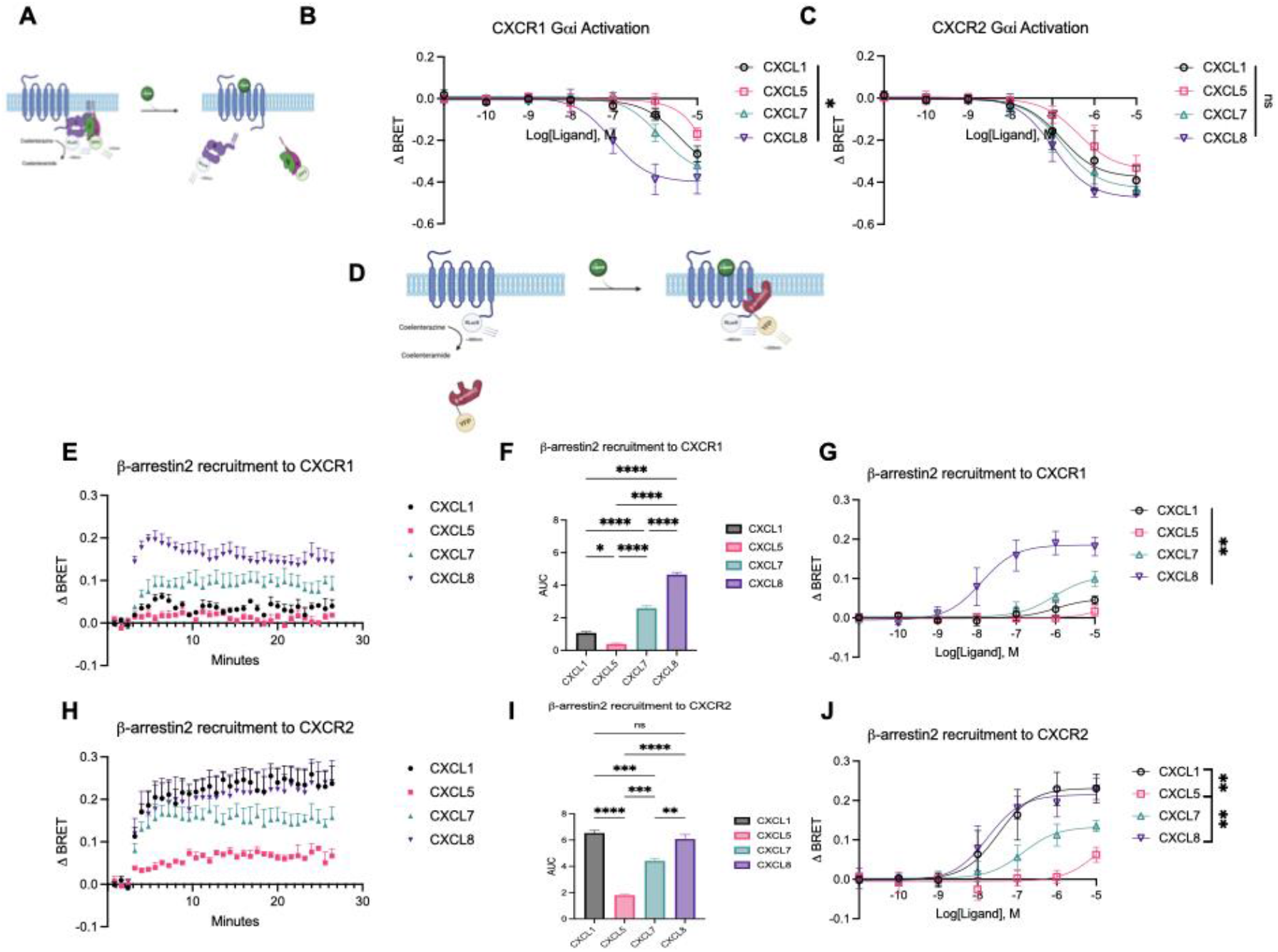
G protein dissociation and β-arrestin 2 recruitment to CXCR1 and CXCR2. **(A)** Schematic of BRET-based TRUPATH assay to assess G protein dissociation. **(B-C)** Dose-response curves of G protein dissociation following receptor activation in HEK293T cells transiently expressing CXCR1 and CXCR2 at listed concentration of chemokine. **(D)** Schematic of BRET-based assay to monitor β-arrestin recruitment to receptor in HEK293T cells transiently expressing β-arrestin2-YFP and either CXCR1-RLuc8 or CXCR2-RLuc8. **(E)** Time course data of β-arrestin2 recruitment to CXCR1 following 10 µM chemokine stimulation **(F)** with corresponding AUC quantification. **(G)** Agonist dose-dependent responses of β-arrestin2 recruitment to CXCR1. **(H)** Time course data of β-arrestin2 recruitment to CXCR2 following 10 μM chemokine stimulation **(I)** with corresponding AUC quantification. **(J)** Agonist dose-dependent responses of β-arrestin2 recruitment to CXCR2. Data for **(B-C)** are measured between 1- and 2-minutes following chemokine addition and are the mean ± SEM, *n* = 3 independent plate-based experiments. Data for **(E-J)** are the mean ± SEM, *n* = 3 independent plate-based experiments. For **(F** and **I)**, a one-way ANOVA followed by Tukey’s multiple comparison’s test was used to compare different chemokines treatments and for **(B-C** and **G-J)** a two-way ANOVA. ns *P* ≥ 0.05, *P<0.05; **P<0.005; ***P<0.0005; ****P<0.0001*, denotes significant effect of ligand by one-way or two-way ANOVA.

### CXCR1 and CXCR2 generate distinct patterns of GRK recruitment to the plasma membrane

We next examined GRK2/GRK3 and GRK5/GRK6 recruitment to the plasma membrane marker CD8α using a previously validated nanoLuc binary technology (nanoBiT) complementation assay (Gardner et al., 2024; Kawakami et al., 2022) (**Fig. 2A**) following stimulation of CXCR1 and CXCR2. Upon activation of CXCR1, chemokine-induced recruitment of GRK2 and GRK3 mirrored the patterns observed in β-arrestin2 recruitment assay, with CXCL8 promoting rapid and maximal recruitment followed by CXCL7 then CXCL1 and CXCL5 (**Fig. 2B** and **F**). A similar pattern was observed with GRK3 (**Fig. 2C** and **G**). Significantly lower recruitment was seen for GRK5 (**Fig. 2D** and **H**) and GRK6 (**Fig. 2E** and **I**), likely due to higher basal signal from kinase already localized to the plasma membrane (Gardner et al., 2024). A similar pattern was seen for GRK5/6 compared to GRK2/3, with more recruitment promoted by CXCL7 and CXCL8 compared to CXCL1 and CXCL5.

**Figure 2.**
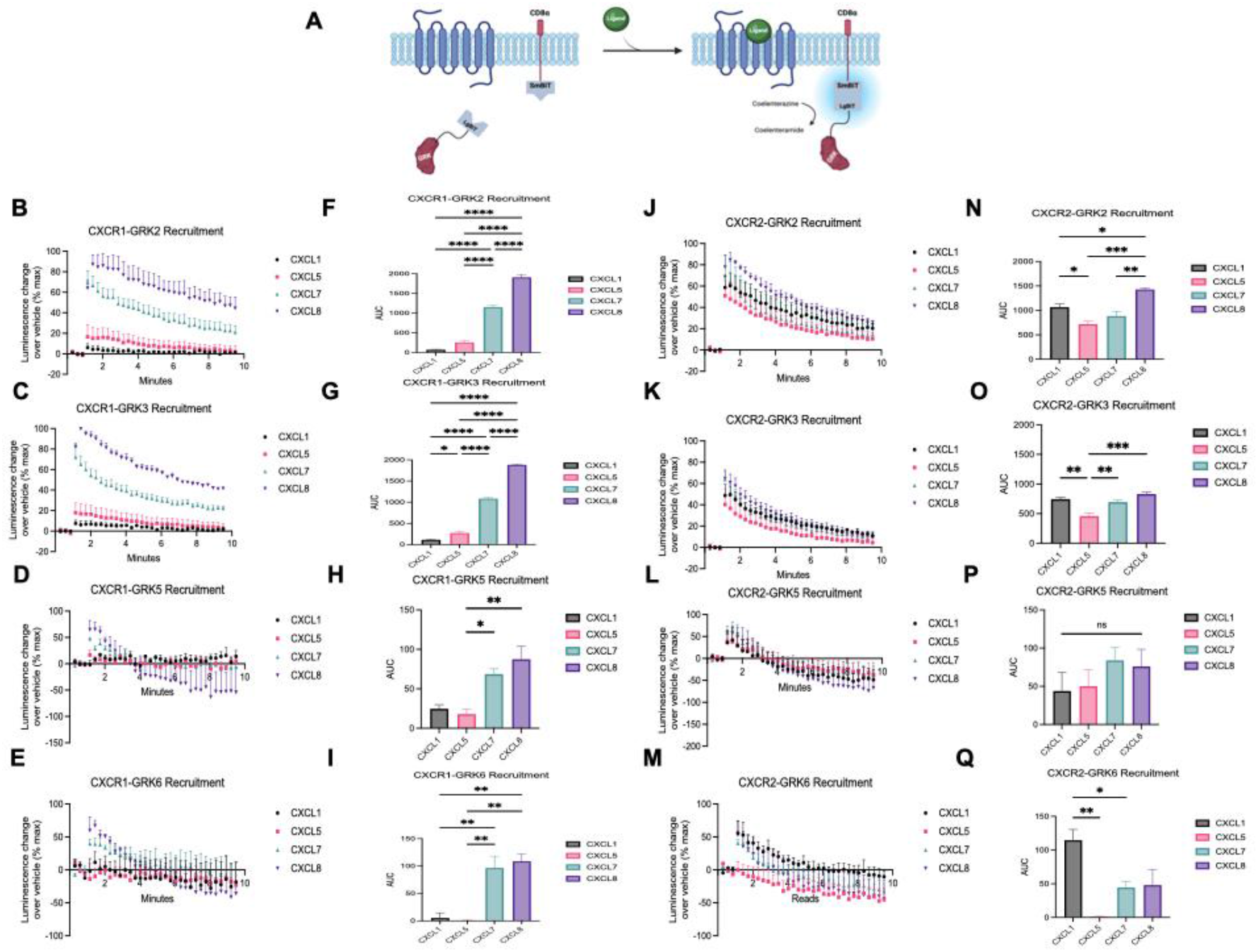
GRK recruitment to plasma membrane following activation of CXCR1 and CXCR2. **(A)** Schematic of NanoBiT luciferase assay to assess GRK recruitment to the plasma membrane using GRK-LgBiT and CD8α-SmBiT constructs in HEK293T cells transiently expressing either CXCR1 or CXCR2. **(B-E)** Time course data of GRK2, GRK3, GRK5, and GRK6 recruitment upon activation of CXCR1 with 10uM of chemokine **(F-I)** with corresponding AUC quantification. **(J-M)** Time course data of GRK2, GRK3, GRK5, and GRK6 recruitment upon activation of CXCR2 with 10 µM of chemokine **(N-Q)** with corresponding AUC quantification. Data for **(B-Q)** are the mean ± SEM, *n* = 3 independent plate-based experiments. For **(F-I and N-Q)** a one-way ANOVA followed by Tukey’s multiple comparison’s test was used to compare different chemokine treatments. ns *P* ≥ 0.05, *P<0.05; **P<0.005; ***P<0.0005; ****P<0.0001*, denotes significant effect of ligand by one-way ANOVA.

At CXCR2, a different trend was observed, with CXCL1 promoting increased GRK2 recruitment along with CXCL7 and CXCL8 (**Fig. 2J** and **N**). Similarly at GRK3, only CXCL5 recruited significantly less GRK3 than the other ligands, with no significant differences observed among the remaining chemokines (**Fig. 2K** and **O**). GRK5 recruitment to the plasma membrane following CXCR2 activation was promoted by all chemokines to similar extents (**Fig. 2L** and **P**) while at GRK6, CXCL1 promoted significantly more recruitment than CXCL7 and 8 (**Fig. 2M** and **Q**).

To evaluate the role of G protein activation in recruiting GRKs to the plasma membrane, we used pertussis toxin (PTX) to inhibit Gαi activation. Following stimulation with CXCL8, we found that PTX significantly reduced both CXCR1- and CXCR2-mediated recruitment of GRK2 and GRK3 (**Supplemental Fig. 2E** and **G**). These findings corroborate previous reports that activity of GRK2/3 family members are largely, but not entirely, dependent on G protein activation due to the presence of a PH domain recognizing free Gβγ following G protein activation (Touhara et al., 1994). Interestingly, GRK6 recruitment following CXCR2 activation was significantly increased upon PTX treatment (**Supplemental Fig. 2H**). Given that GRK6 is constitutively localized to the plasma membrane (Loudon and Benovic, 1997), this apparent increase may have been the result of compensatory recruitment following a decrease of GRK2/3 recruitment in response to G protein inhibition.

### Chemokines promote different amounts of CXCR1 and CXCR2 internalization

To determine the functional relevance of the trends observed in β-arrestin2 and GRK recruitment, we next assessed agonist-induced internalization of CXCR1 and CXCR2. Receptor internalization was monitored by BRET between luciferase-tagged CXCR1 and CXCR2 and FYVE domain-tagged mVenus in endosomes (**Fig. 3A**). At CXCR1, rank order of internalization mirrored the trends observed in β-arrestin and GRK2/3 recruitment-CXCL8 > CXCL7 > CXCL1 > CXCL5 (**Fig. 3B-D**). However at CXCR2, the differences across all chemokines except CXCL5 was statistically insignificant. CXCL5 promoted significantly less internalization (**Fig. 3E-G**), a trend consistent with the pattern of β-arrestin2 recruitment to CXCR2 (**Fig. 1I**).

**Figure 3.**
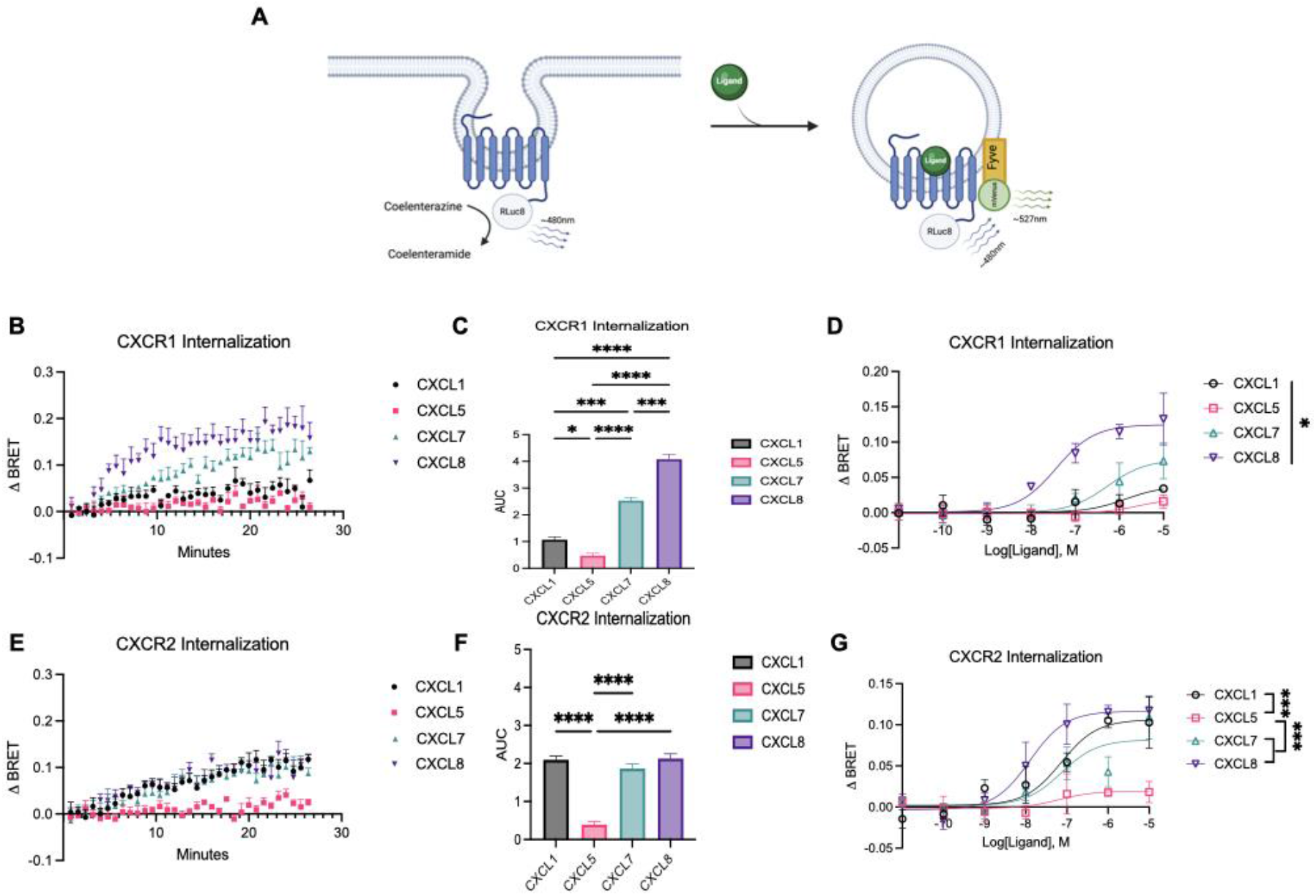
CXCR1 and CXCR2 internalization. **(A)** Schematic of BRET-based assay to monitor receptor trafficking to early endosomes using the BRET acceptor Fyve-mVenus in HEK293T cells transiently expressing either CXCR1-RLuc8 or CXCR2-RLuc8. **(B)** Time course data of CXCR1-RLuc8 trafficking to early endosomes following stimulation with 10 µM of chemokine **(C)** with corresponding AUC quantification. **(D)** Agonist dose-dependent responses of CXCR1-RLuc8 internalization between 10- and 11-minutes after ligand addition and are the mean ± SEM, *n* = 3 independent plate-based experiments. **(E)** Time course data of CXCR2-RLuc8 trafficking to early endosomes following stimulation with 10 µM of chemokine **(F)** with corresponding AUC quantification. **(G)** Agonist dose-dependent responses of CXCR2-RLuc8 internalization between 25- and 26-minutes after ligand addition and are the mean ± SEM, *n* = 3 independent plate-based experiments. For **(C and F)**, a one-way ANOVA followed by Tukey’s multiple comparison’s test was used to compare different chemokines treatments and for **(D and G)** a two-way ANOVA. ns *P* ≥ 0.05, *P<0.05; **P<0.005; ***P<0.0005; ****P<0.0001*, denotes significant effect of ligand by one-way or two-way ANOVA.

Given these observations, we sought to determine the role of β-arrestins in promoting internalization of CXCR1 and CXCR2 following treatment with CXCL8 in β-arrestin 1/2 KO cells. Compared to our findings in HEK293T cells, internalization of both receptors was significantly reduced, but not completely abrogated (**Supplemental Fig. 3A** and **B**). To validate receptor internalization occurs independent of G protein activation, we repeated these internalization studies in β-arrestin 1/2 KO cells treated with PTX; no differences in CXCR1 and CXCR2 internalization upon stimulation with CXCL8 were observed (**Supplemental Fig. 3C** and **D**). Together, our findings suggest that chemokine-induced internalization of CXCR1 and CXCR2 is primarily mediated by β-arrestins.

### Assessment of ligand bias at CXCR1 and CXCR2

To perform a qualitative analysis of bias (Rajagopal et al., 2011), we constructed “bias plots” to identify biased agonists for each chemokine at CXCR1 and CXCR2. Bias plots allow for an assessment of assay amplification by converting dose-response data for two signaling pathways of interest and plotting the responses of these two different assays at the same concentration of ligand against each other. The resulting curve shape allows for a direct comparison of signaling through the two different pathways that varies depending on the sensitivities of the assays being compared; thus “test” agonists must be qualitatively compared to a balanced reference agonist (Rajagopal et al., 2011). We compared G protein dissociation, β-arrestin1 and β-arrestin2 recruitment, and receptor internalization. Using CXCL8 as the reference balanced agonist, as it robustly signals through both G protein and β-arrestins, CXCL7 demonstrated a relative decrease in both G protein activation and β-arrestin coupling and internalization, acting as a partial biased agonist at CXCR1. Conversely, CXCL1 and CXCL5 displayed relative bias towards G protein activation compared to CXCL7 and CXCL8 (**Fig. 4A-C**). At CXCR2, we observed a shift in these trends. While CXCL7 and CXCL8 acted as balanced agonists and CXCL5 as a G protein-biased agonist, CXCL1 demonstrated an appreciable shift towards β-arrestin coupling and internalization while preserving its ability to activate G protein (**Fig. 4D-E**). Although CXCL1 is a relatively G protein-biased partial agonist at CXCR1, it behaves as a balanced full agonist similar to CXCL8 at CXCR2.

**Figure 4.**
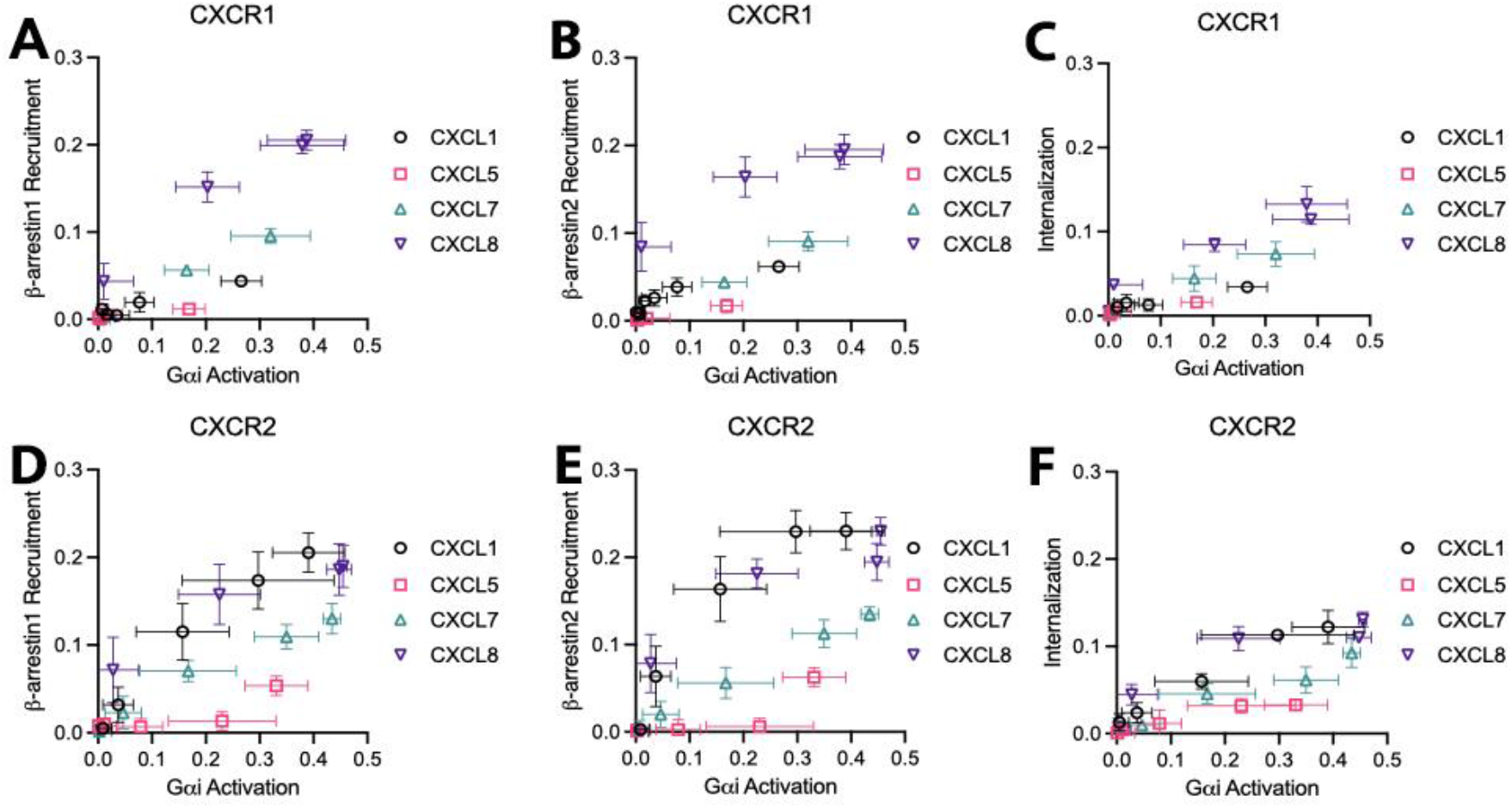
Bias plots to qualitatively assess relative bias across signaling pathways at CXCR1 and CXCR2. **(A)** β-arrestin1 recruitment and Gαi activation; **(B)** β-arrestin2 recruitment and Gαi activation; **(C)** internalization and Gαi activation. CXCR2 bias plots of **(D)** β-arrestin1 recruitment and Gαi activation; **(E)** β-arrestin2 recruitment and Gαi activation; **(F)** internalization and Gαi activation.

### CXCR1 and CXCR2 activate distinct subcellular pools of ERK

Lastly, to assess distal signaling downstream of CXCR1 and CXCR2, we examined activation of the mitogen activated-protein kinase (MAPK) pathway through ERK1/2 phosphorylation (pERK) at different subcellular locations using previously validated BRET-based ERK activity reporter biosensors (EKAR) targeted to the cytoplasm, nucleus, and early endosomes (Pham et al., 2024) (**Fig. 5A**). We found that chemokines induced unique location-specific ERK activity following CXCR1 activation. CXCL7 promoted more ERK activation than CXCL5 in the cytoplasm (**Fig. 5B** and **E**), while CXCL8 activated more ERK than CXCL1 and CXCL5 in the nucleus (**Fig. 5D** and **G**). Although all chemokines promoted endosomal ERK activity, no statistically significant differences were observed amongst the chemokines (**Fig. 5C** and **F**). Interestingly, while CXCR2 activated ERK in the cytoplasm (**Fig. 5H** and **K**), nucleus (**Fig. 5J** and **M**), and endosomes (**Fig. 5**Iand **L**), no statistically significant differences were observed across chemokines. To further dissect the role of β-arrestin-mediated internalization and G protein activation in promoting ERK activity, we inhibited receptor-internalization using a dominant-negative mutant of the GTPase Dynamin (Dynamin K44A), which is required for clathrin-mediated endocytosis (Damke et al., 1994), along with PTX to inhibit G protein activation. Following treatment of CXCR1 and CXCR2 with the full endogenous agonist CXCL8, PTX treatment alone significantly decreased endosomal ERK activity compared to pcDNA control at CXCR1 (**Supplemental Fig. 4A-D**). However, inhibition of internalization alone had minimal effect on endosomal ERK activity at CXCR1 and CXCR2. Simultaneous inhibition of both G proteins and internalization led to an additive decrease in ERK activation at both receptors, suggesting both β-arrestin-mediated internalization and G protein activation are required for maximal endosomal ERK activity at CXCR2, whereas G protein activation primarily mediates endosomal ERK activity at CXCR1. These observations validate previous findings (Prado et al., 2007) which suggested CXCR1 stabilizes a receptor signaling state at the plasma membrane, while CXCR2 promotes sustained ERK activation via G protein-dependent and G protein-independent modes at the plasma membrane and following internalization, respectively.

**Figure 5.**
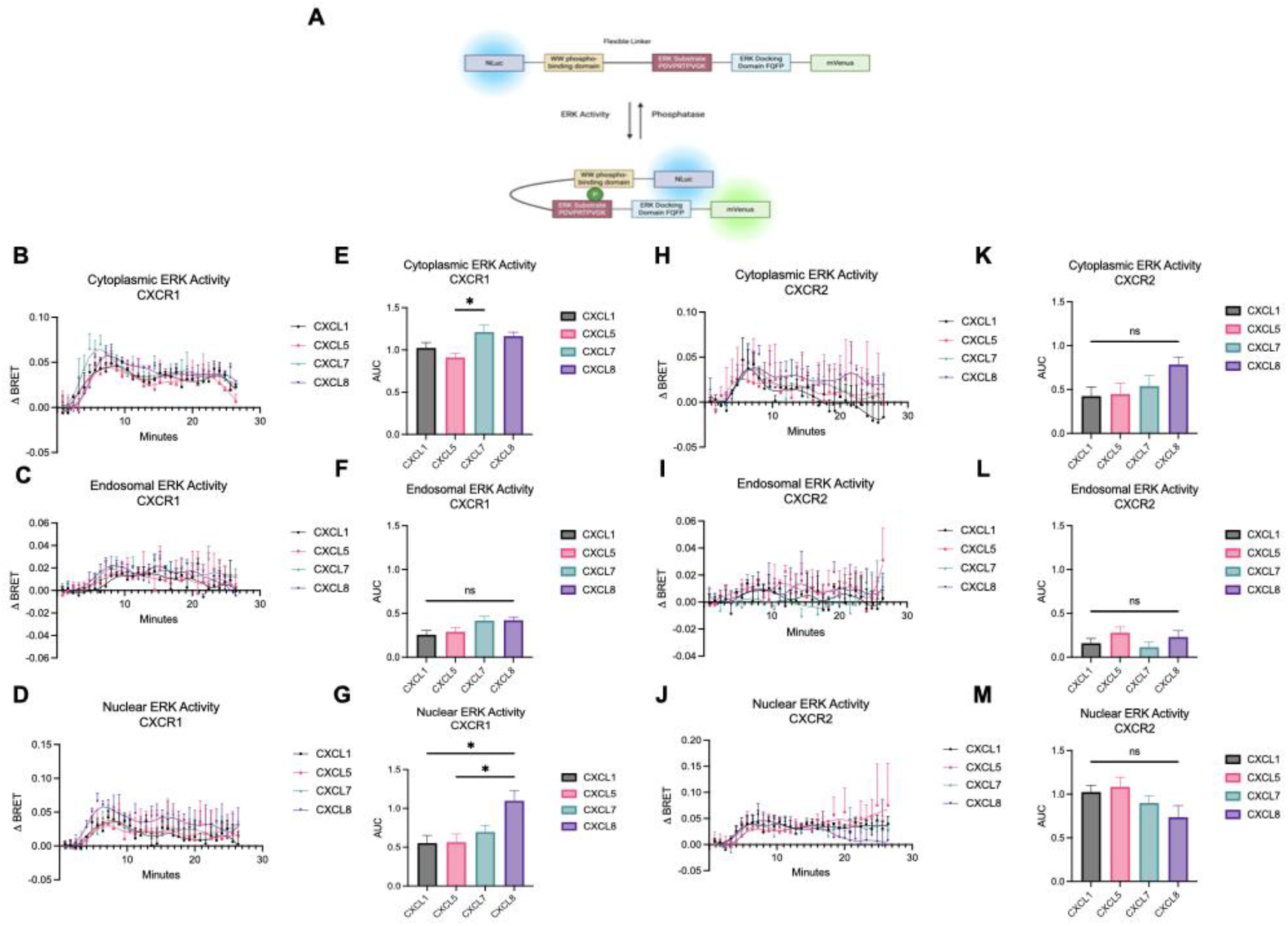
CXCR1 and CXCR2 promote distinct ERK activation patterns at different subcellular locations. **(A)** Schematic of subcellular EKAR BRET biosensors targeted to the cytoplasm, the nucleus, and early endosomes to measure location-specific ERK activity induced by endogenous chemokines of CXCR1 and CXCR2 in HEK293T cells. **(B-D)** Time course data of cytoplasmic, nuclear, and endosomal ERK activity following stimulation of CXCR1 with 10 µM of chemokine **(E-G)** with corresponding AUC quantification. **(H-J)** Time course data of cytoplasmic, nuclear, and endosomal ERK activity following activation of CXCR2 with 10 µM of chemokine **(K-M)** with corresponding AUC quantification. Data for **(B** and **E)** are the mean ± SEM, *n* = 4 independent plate-based experiments and **(C-D; F-M)** are the mean ± SEM, *n* = 3 independent plate-based experiments. For **(E-G** and **K-M)**, a one-way ANOVA followed by Tukey’s multiple comparison’s test was used to compare different chemokines treatments; ns *P* ≥ 0.05, *P<0.05; **P<0.005; ***P<0.0005; ****P<0.0001*, denotes significant effect of ligand by one-way ANOVA.

## DISCUSSION

In this study, we observe that the same chemokine agonists for CXCR1 and CXCR2 display different patterns of transducer and effector activation. ELR-chemokines play diverse roles including in normal homeostasis to combating bacterial infections in most organs, and are also selectively expressed to different insults in different organs (Rajarathnam et al., 2019). For instance, CXCL8 is preferentially expressed in lung infections, and CXCL7 is highly expressed and released from platelets regulating neutrophil-platelet crosstalk and thrombosis-related diseases. The receptors are predominantly expressed in neutrophils but also in non-immune cells. Our current study characterizing the activity of proximal transducers suggest that chemokines are not redundant, their functional phenotype is unique, and that detailed knowledge of receptor-specific signalizing activities mediated by intracellular signaling machinery in conjunction with their expression levels and tissue locations are essential to fully understand the role of signaling bias in determining chemokines’ role in physiology and pathology.

The most notable difference was the pattern of signaling by CXCL1, which demonstrated clear evidence of receptor bias between CXCR1 and CXCR2. At CXCR1, CXCL8 robustly activated more Gαi than CXCL1, CXCL5, and CXCL7. Similarly, in rank order, CXCL8 followed by CXCL7, CXCL1, and CXCL5 recruited both β-arrestin1 and β-arrestin2. Interestingly, a different pattern was observed at CXCR2. At CXCR2, CXCL1 demonstrated β-arrestin recruitment pattern similar to CXCL8. At both receptors, the patterns of receptor internalization mirrored their patterns of β-arrestin recruitment. It is important to note that β-arrestins are largely, but not entirely, responsible for mediating receptor internalization as shown in our studies using β-arrestin 1/2 KO cells. Using bias plots to assess these chemokine signaling profiles, we found that CXCL1 behaves as a G protein-biased partial agonist similar to CXCL5 and CXCL7 at CXCR1, but as a balanced full agonist similar to CXCL8 at CXCR2. Our findings at CXCR1 are consistent with a recent report (Boon et al., 2025) that identified CXCL7, a high-affinity ligand, and CXCL1 and CXCL5, low-affinity ligands, as G protein-biased compared to the reference ligand CXCL8. Notably, previous studies investigating ligand bias at CXCR2 concluded CXCL1 to be G protein-biased when compared to CXCL8 (Boon et al., 2024; Rajagopal et al., 2013), whereas we found CXCL1 behaved as a balanced agonist similar to CXLC8. It is important to note bias plots are not without limitations, since factors such as systems bias may still qualitatively change relative differences observed between pathways.

As CXCR1 and CXCR2 predominantly couple to GRK2 and GRK6, respectively, to regulate neutrophil-mediated inflammatory responses (Raghuwanshi et al., 2012), we focused our studies on the recruitment of the GRK2/3 and GRK5/6 families to both receptors. We found that the rank order of chemokine-induced recruitment of GRK2 and GRK3 were similar to the trends observed in β-arrestin recruitment to CXCR1. However, the rank order of chemokine-mediated GRK recruitment following CXCR2 activation varied across GRKs with recruitment of no single GRK correlating directly with β-arrestin recruitment or receptor internalization. Our data suggests that the interplay of receptor and chemokine bias at CXCR1 and CXCR2 may differentially modulate the functional output of unique GRK ensembles to produce unique phosphorylation barcodes at these receptors, as shown in previous studies at the chemokine receptor CXCR3 (Eiger et al., 2023). However, further investigation is necessary to directly test this hypothesis by mapping the phosphorylation of specific sites to individual kinases.

Like many GPCRs, CXCR1 and CXCR2 have been shown to engage the MAPK pathway following treatment with CXCL8 (Nasser et al., 2009; Prado et al., 2007; Raghuwanshi et al., 2012). We expand on these findings to include CXCL1, CXCL5, and CXCL7, further illuminating that these chemokines promote similar subcellular pools of ERK within the cytoplasm, nucleus, and early endosomes. Moreover, it was appreciated that both G protein activation and receptor internalization contribute to promoting endosomal ERK for both CXCR1 and CXCR2. These findings enhance the granularity of signaling activity of CXCR1 and CXCR2 and their endogenous agonists through the lens of location-bias. Further studies are needed to assess the specific function of these unique signaling profiles in the context of mediating neutrophil activity during inflammatory responses.

## Nonstandard abbreviations

GPCR: G protein-coupled receptor
GRK: G protein receptor kinase
CKR: chemokine receptor
BRET: bioluminescence resonance energy transfer
RLuc: renilla luciferase
NLuc: NanoLuc luciferase
EKAR: ERK activity reporter biosensors
NanoBiT: nanoLuc binary technology
PTX: pertussis toxin

## ACKNOWLEDGEMENT

We thank Nour Nazo for laboratory assistance. SR is funded by R01 GM122798 and KR by NIH grants R21 AI160613 and a pilot grant from the Institute for Human Infections and Immunity. Diagrams were made using BioRender.

---

**Supplemental Figure 1.**
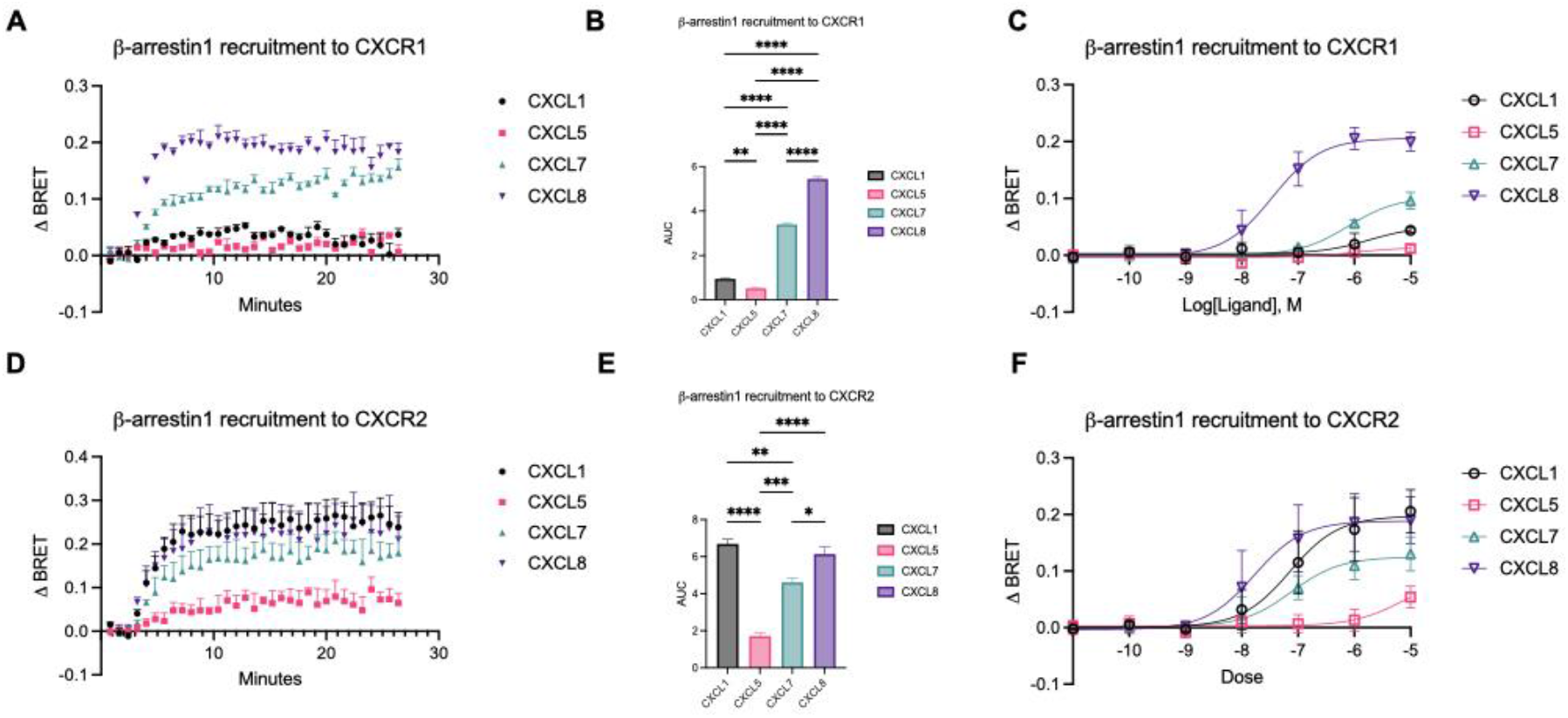
HEK293T cells transiently expressing β-arrestin2-YFP and either CXCR1-RLuc8 or CXCR2-RLuc8 were treated with 10 µM of chemokine. **(A)** Time course data of β-arrestin1 recruitment to CXCR1 **(B)** with corresponding AUC quantification. **(C)** Agonist dose-dependent responses of β-arrestin1 recruitment to CXCR1. **(D)** Time course data of β-arrestin1 recruitment to CXCR2 following 10 μM chemokine stimulation **(E)** with corresponding AUC quantification. **(F)** Agonist dose-dependent responses of β-arrestin1 recruitment to CXCR2. Data for **(C and F)** are measured between 1- and 2-minutes following chemokine addition. Data for **(A-E)** are the mean ± SEM, *n* = 3 independent plate-based experiments. For **(B and E)**, a one-way ANOVA followed by Tukey’s multiple comparison’s test was used to compare different chemokines treatments and for **(C and F)** a two-way ANOVA. ns *P* ≥ 0.05, *P<0.05; **P<0.005; ***P<0.0005; ****P<0.0001*, denotes significant effect of ligand by one-way or two-way ANOVA.

**Supplemental Figure 2.**
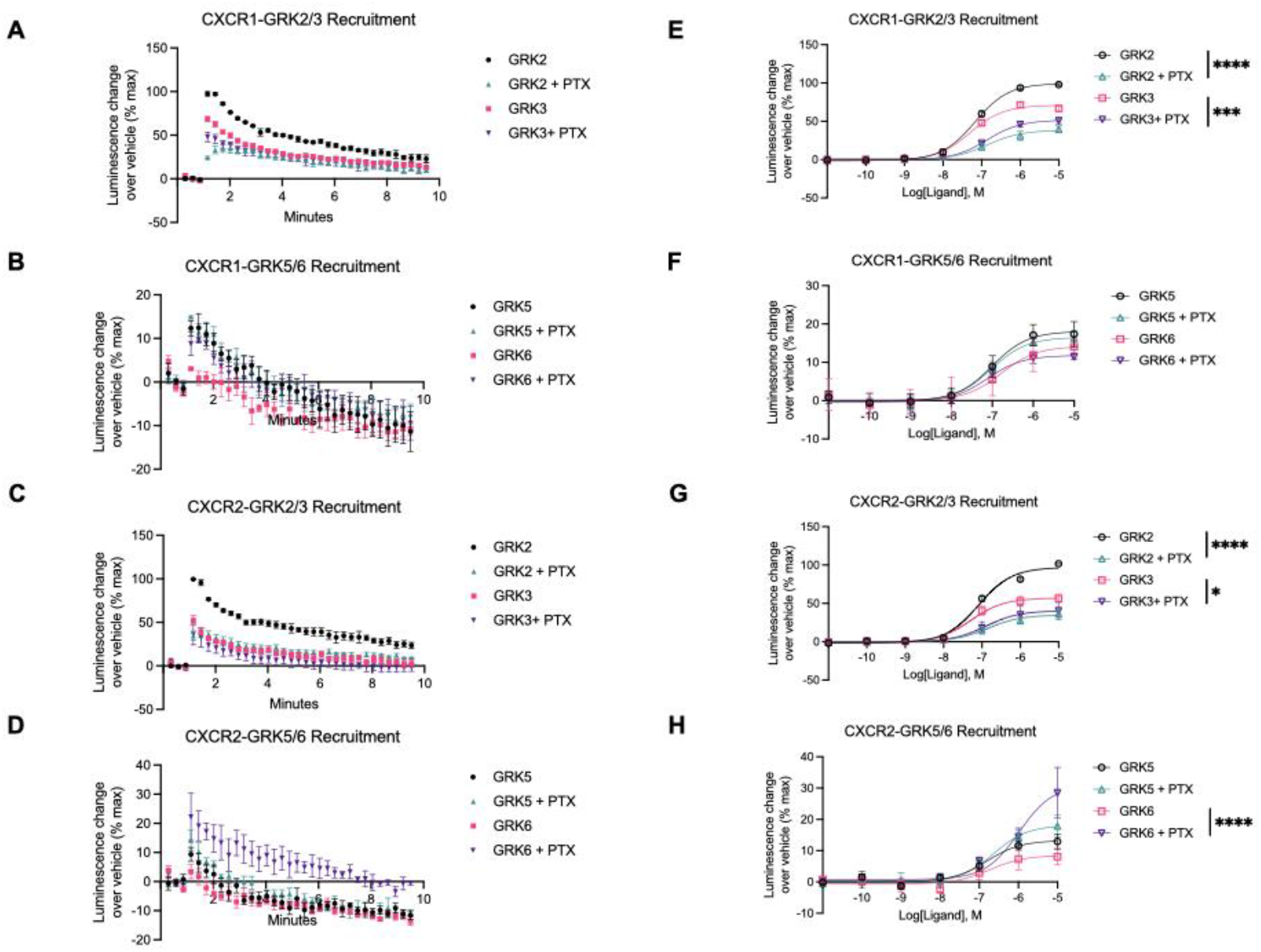
NanoBiT complementation assay to assess GRK recruitment to the plasma membrane using GRK-LgBiT and CD8α-SmBiT constructs in HEK293T cells transiently expressing either CXCR1 or CXCR2. **(A)** Kinetic time-course data of GRK2/GRK3 and **(B)** GRK5/6 recruitment upon activation of CXCR1 using 10µM of CXCL8 with and without pretreatment of 200ng/mL pertussis toxin **(E and F)** with corresponding dose-response curves. **(C)** Kinetic time-course data of GRK2/GRK3 and **(D)** GRK5/6 recruitment upon activation of CXCR2 using 10µM of CXCL8 with and without pretreatment of 200ng/mL pertussis toxin **(G and H)** with corresponding dose-response curves. Data for **(A-H)** are the mean ± SEM, *n* = 3 independent plate-based experiments; **(E-H)** A two-way ANOVA of ligand and concentration was performed. Ns *P* ≥ 0.05, *P<0.05; **P<0.005; ***P<0.0005; ****P<0.0001*.

**Supplemental Figure 3.**
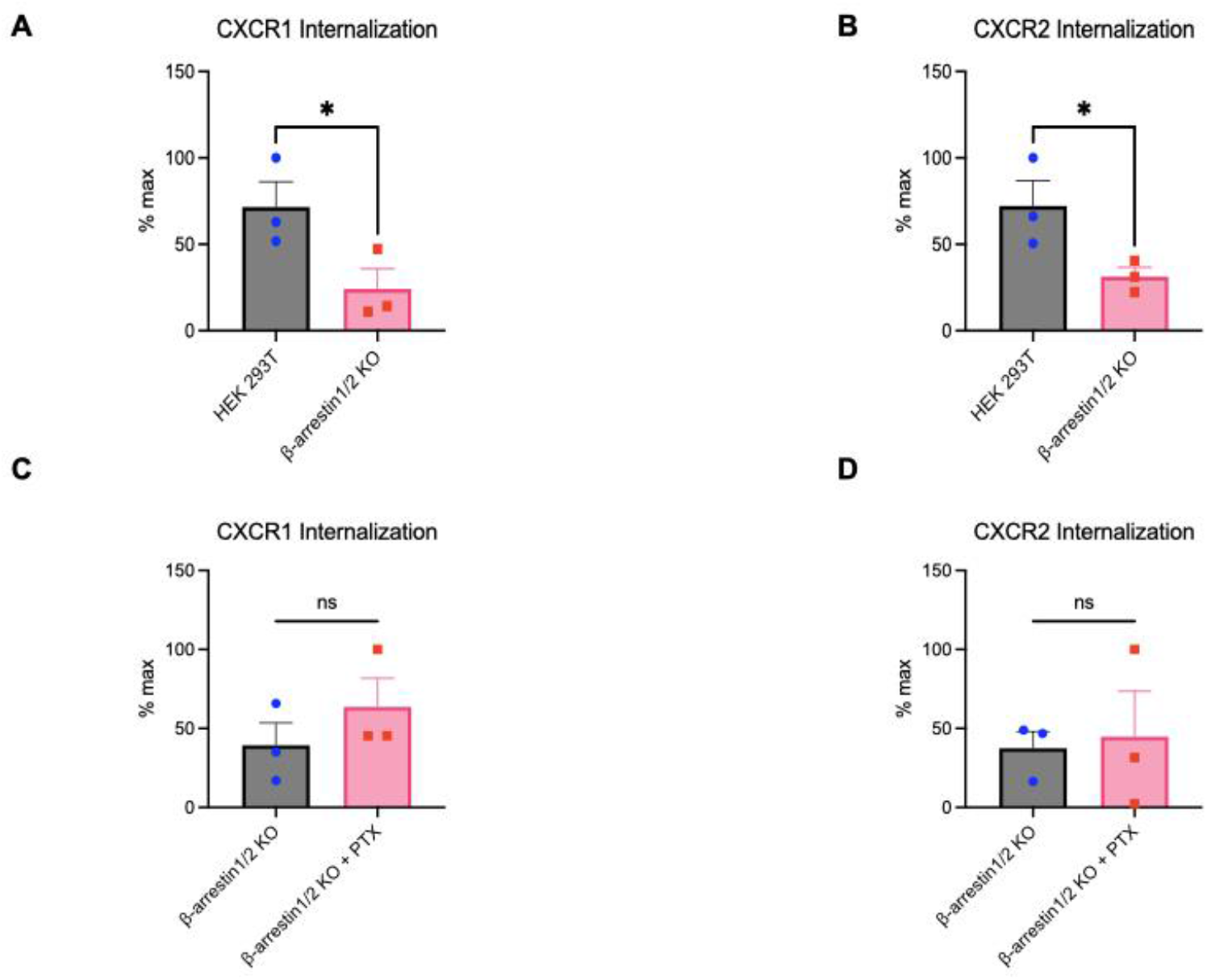
BRET-based assay to monitor receptor trafficking to early endosomes using the BRET acceptor Fyve-mVenus in HEK293T and β-arrestin1/2 KO cells transiently expressing either CXCR1-RLuc8 or CXCR2-RLuc8. **(A)** CXCR1-RLuc8 and **(B)** CXCR2-RLuc8 internalization response comparisons in HEK293T vs β-arrestin1/2 KO cells which are normalized to maximum signal measured between 20- and 21-min following 10 µM CXCL8 treatment. **(C)** CXCR1-RLuc8 and **(D)** CXCR2-RLuc8 internalization responses in β-arrestin1/2 KO cells with or without pretreatment of 200 ng/mL pertussis toxin normalized to maximum signal measured between 4- and 5-min following 10 µM CXCL8 treatment. Data for **(A-D)** are the mean ± SEM, *n* = 3 independent plate-based experiments. An unpaired t-test was used for **(A-D)**. ns *P* ≥ 0.05, *P<0.05; **P<0.005; ***P<0.0005; ****P<0.0001*.

**Supplemental Figure 4.**
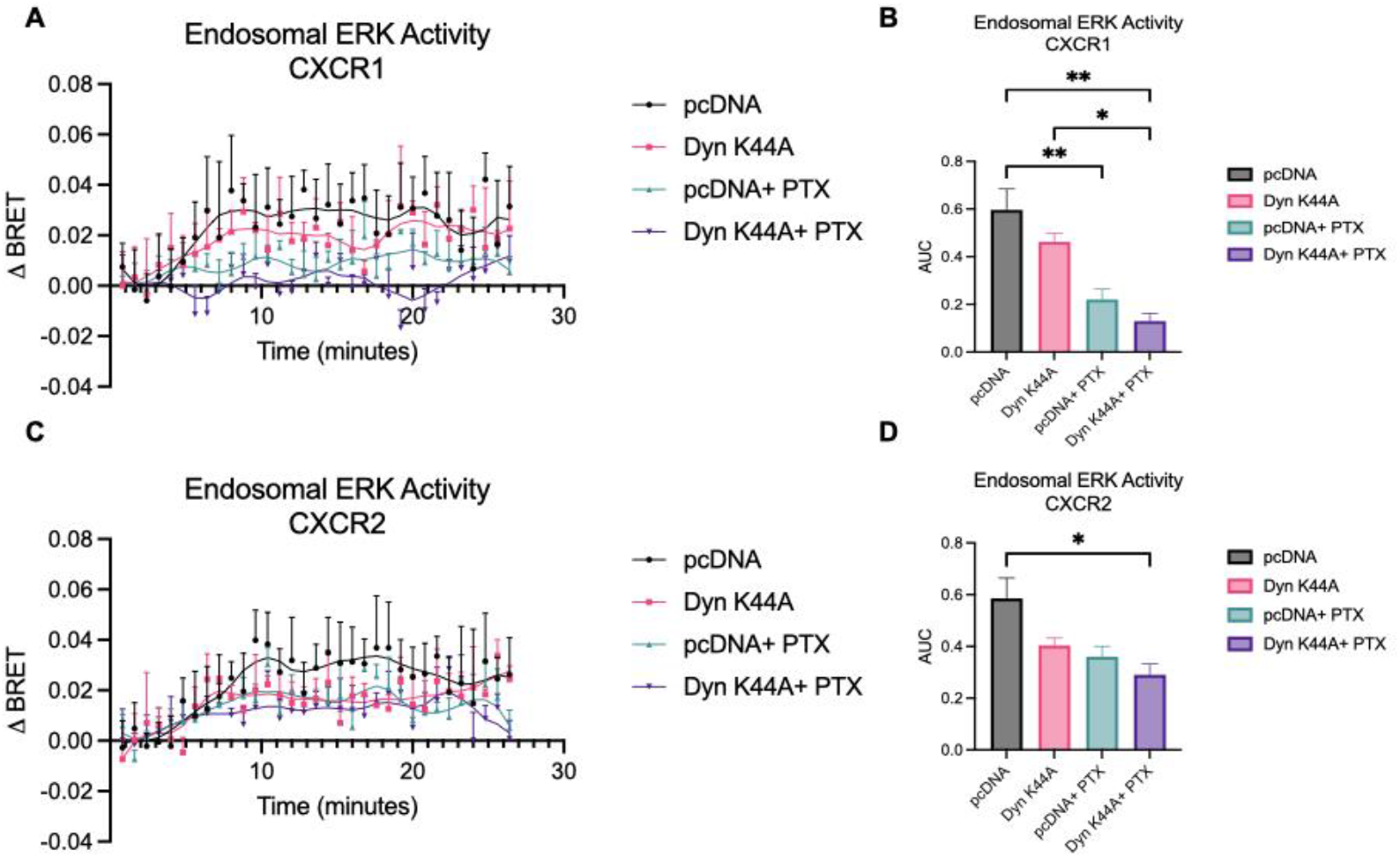
**(A)** Kinetic time-course data of endosomal ERK activity in HEK293T cells transiently expressing pcDNA or Dynamin K44A with or without pretreatment of 200ng/mL pertussis toxin following stimulation of CXCR1 using 10µM of CXCL8 **(B)** with corresponding AUC quantification. **(C)** Kinetic time-course data of endosomal ERK activity in HEK293T cells transiently expressing pcDNA or Dynamin K44A with or without pretreatment of 200ng/mL pertussis toxin following stimulation of CXCR1 with 10µM of CXCL8 **(D)** with corresponding AUC quantification. For **(B and D)**, a one-way ANOVA followed by Tukey’s multiple comparison’s test was used to compare different treatment conditions; ns *P* ≥ 0.05, *P<0.05; **P<0.005; ***P<0.0005; ****P<0.0001*, denotes significant effect of ligand by one-way ANOVA.

**Supplementary Table 1:**
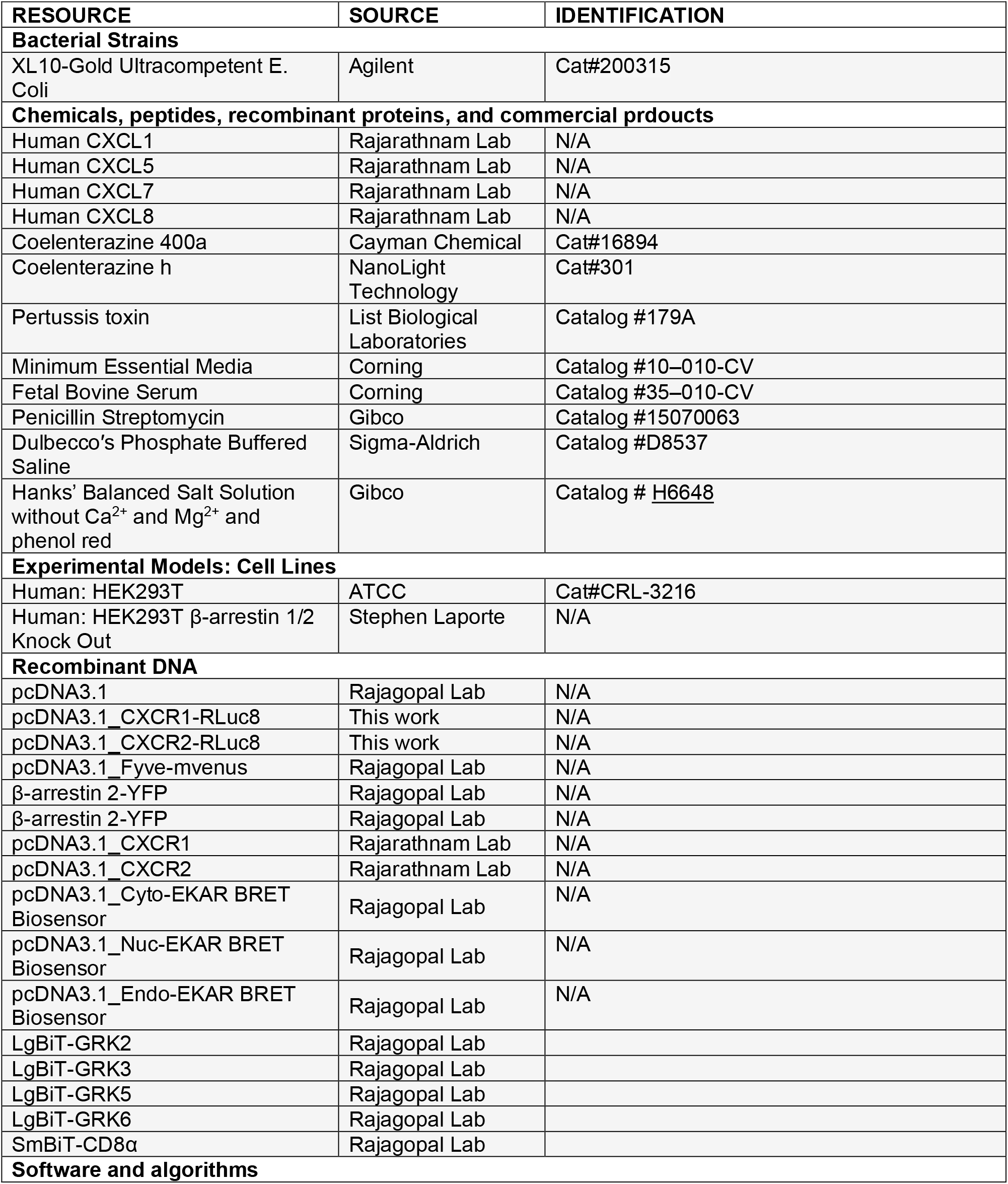
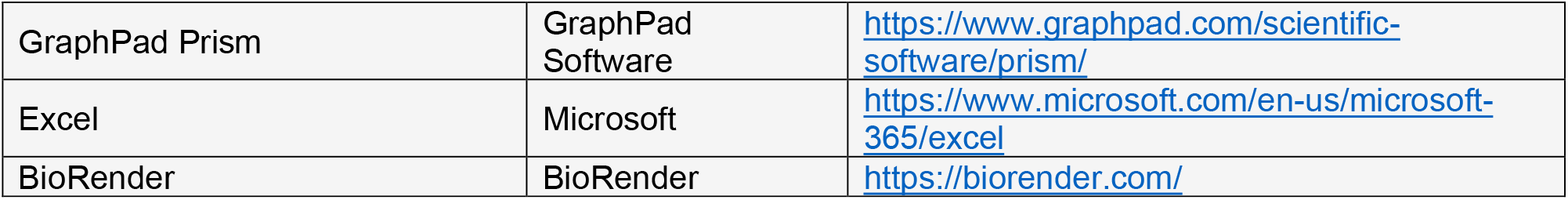
Key resources and DNA constructs used in this study.

